# The Mangrove Microbiome of the Malay Peninsula

**DOI:** 10.1101/2022.11.21.517418

**Authors:** Benjamin J. Wainwright, Trevor Millar, Lacee Bowen, Kate Hickman, Jen Nie Lee, Zee Yi Yeo, Danwei Huang, Geoffrey Zahn

## Abstract

Microbes have fundamental roles underpinning the functioning of our planet, they are involved in global carbon and nutrient cycling, and support the existence of multicellular life. The mangrove ecosystem is nutrient limited and without microbial cycling, life in this harsh environment would likely not exist. The mangroves of Southeast Asia are the oldest and most biodiverse of all the planets. They have vital roles helping to prevent shoreline erosion, act as nursery grounds for many marine species and contain significant stocks of sequestered carbon. Despite these recognised benefits and the importance of microbes in these ecosystems, studies examining the mangrove microbiome are scarce, especially in the Southeast Asian biodiversity hotspot. Here we examine the microbiome of *Avicenia alba* and *Sonneratia alba* and identify a core microbiome of 81 taxa, a further eight taxa (*Pleurocapsa, Tunicatimonas, Halomonas, Marinomonas, Rubrivirga, Altererythrobacte, Lewinella*, and *Erythrobacter*) were found to be differentially abundant suggesting key roles in this microbiome, with the identified dimethylsulfoniopropionate (DMSP) metabolisers having important functions in these habitats. The majority of those identified are involved in nutrient cycling or involved in the production of compounds that promote host survival. Increasingly, blue carbon and nature-based solutions to climate change are heralded as viable mitigation steps to limit climate change, however, this is done with little to no consideration of the microbial communities that cycle sequestered carbon in these environments. Here, we examine the microbial communities present in sediment samples taken in close proximity to each tree, sediment samples represent a major sink of atmospheric carbon and understanding how the associated communities will change as climate change advances will become an increasingly important part of carbon stock assessments. Knowing what microbes are presently there is an important first step in this process.

## Introduction

Mangrove trees occupy a transitional zone between marine and terrestrial environments, and are the only tree species on the planet that can thrive in the saline, oxygen limited environment found in this transition zone [1–4]. These highly productive ecosystems are found in tropical and subtropical regions where they have significant ecological and economic importance. They are critical for nutrient cycling, providing basal organic matter to coastal food webs, preventing coastal erosion, filtering pollutants, are biodiversity hotspots, and act as nurseries for many marine animals [5–8]. Economically, they buffer against natural disasters, support coastal fisheries, and are the source of myriad forestry products [8, 9].

Microorganisms have key roles in mangrove ecosystems, and are important in promoting growth and maintaining productivity [10, 11]. These microbes contribute significantly to carbon cycling and the global carbon budget. Despite their limited area, accounting for only approximately 3% of global forest cover, mangroves are significant carbon sinks with estimates suggesting they contain 10% of all global carbon emissions [12–14]. Known as “blue carbon”, mangroves are able to sequester atmospheric carbon dioxide in above and below ground structures (e.g., leaves and roots etc). Ultimately, this carbon becomes locked in the anoxic sediments where it remains stable until disturbed [14, 15].

Once disturbed, usually through anthropogenic factors such as habitat clearance or modification, microbial processes can release this carbon back into the atmosphere where it contributes to climate change [16, 17]. However, despite the growing interest in blue carbon, we currently lack a detailed understanding or characterisation of the microbial communities and potential drivers involved in biogeochemical cycling in these coastal ecosystems [18], particularly in Southeast Asia. As climate change progresses and coastal habitats are further degraded by land use changes, eutrophication, and other anthropogenic activities, many of these coastal ecosystems are predicted to become net sources of carbon instead of sinks as microbial communities change in response and sequestered carbon becomes available for microbial cycling [17, 19–24]. Consequently, mangroves and other coastal habitats, along with their associated blue carbon stocks, are unlikely to fulfil their claimed potential as nature based climate solutions, and many of their benefits are likely overstated [15, 25–28]

The importance of the microbiome in maintaining and promoting host health has been recognised in many organisms in both terrestrial and marine environments [29–33], and the role of the microbiome in promoting plant growth and resilience in terrestrial plants is well established [34, 35]. However, the ecological importance of the coastal microbiome is not yet well-understood, although several studies have shown that fungal and bacterial diversity are key to habitat restoration in marine environments [5, 8]. Several mangrove-associated bacteria are known to promote root growth, support nutrient cycling and availability, degrade contaminants, and aid in other essential processes [7, 36, 37]. Despite this recognised importance, the dynamics, distribution, and community composition of microbes in mangrove ecosystemsremains vague [10, 11, 38]. This paucity is particularly acute in Southeast Asian coastal regions, but see [39, 40] for relevant work examining seagrass ecosystems.

Thirty four percent of global mangrove cover is found in Southeast Asia, with Indonesia and Malaysia having the largest area of mangrove forest in the region [41–43]. Mangroves in Southeast Asia are typically highly productive, and are the oldest and most biodiverse mangrove forests in the world [44–46]. Yet, primarily a consequence of anthropogenic activities [47–49], regional rates of loss are some of the highest in the world [43]. Restoration of mangrove ecosystems through strategies such as out-planting of nursery-raised saplings, raised bed methods, and direct propagule planting have shown mixed results, with most eventually failing [46, 50–53]. However, greater success has been achieved when inoculation with local bacterial and fungal species has been performed [5] and the matching of microbial communities between transplant and out-planting sites to avoid host maladaptation to a new environment has been recommended [54, 55]. Given that microbial communities frequently show differences in composition, even across comparatively small spatial scales, an understanding of this community structure should be an important consideration in future restoration activities [56–60].

The idea of a ‘core microbiome’ was initially explored by the Human Microbiome Project, and was defined as a group of microbial taxa that are shared by all, or most humans [61]. The ubiquity of these taxa in their host species led to the suggestion that this core microbiome may play an important role in maintaining host biological function [62], here we examine the microbiome of two widespread mangrove species, *Sonneratia alba* and *Avicennia alba* throughout the Malay Peninsula. We hypothesise that an overall ‘core mangrove microbiome’ will exist and, at a smaller scale, a core microbiome will be shared between different tree structures (e.g., leaves vs pneumatophores) within species and between species. More generally, despite sharing a core taxa of microbes throughout the region and between species we expect to identify microbial community differentiation between species, sample sites, and sampled structure.

Studies such as this are an important step in understanding the Southeast Asian mangrove microbiome, and they allow us to generate baseline data on the associated microbial communities. These communities are predicted to change under future projected climate change scenarios [63], but it is impossible to assess the magnitude of these changes, or the taxonomic shifts that will occur without knowing what is presently there and in which locations.

## Methods

At each of nine sampling locations throughout Singapore and Malaysia we targeted 10 visibly healthy whole leaves, fruiting bodies (mangrove fruit) and entire pneumatophores from *Anicennia alba* and *Sonneratia alba*. In addition to living tissue, a sediment sample was collected in close proximity to each tree (<1m), with sediment samples taken from approximately 4cm below the surface. We were unable to find *Anicennia alba* at two sample sites in Malaysia (Port Dickson, and Tioman) (Fig. 1). For both species, DNA was extracted with a Qiagen DNeasy PowerSoil Kit following the manufacturer’s protocol. Prior to extraction all samples were disrupted in an Omni Bead Ruptor 24 (Omni International, Kennesaw, GA, United States) at 8 ms^− 1^ for 2min. PCR amplification targeting the V4 region of the 16S SSU rRNA gene was performed using the 515F and 806R primers modified to include Illumina adaptors, a linker and a unique barcode [64]. All reactions were performed in a total volume of 25 µl, containing 1 µl of undiluted template, 0.1 µl of KAPA 3G Enzyme (Kapa Biosystems, Inc, Wilmington, MA, USA), 0.75 µl of each primer at 10 µM, 2.5 µl, 1.5 µl of 1.5 mg ml^-1^ BSA, 12.5 µl KAPA PCR Buffer and water to 25 µl. PCR cycling was 94°C for 180 s, followed by 35 cycles of 94°C for 45 s, 50°C for 60 s and 72°C for 90 s, and a final extension at 72°C for 10 min. Negative extraction and PCR controls were included to identify any potential contamination issues. Prior to pooling for sequencing, normalisation and cleaning of PCR products was performed in SequalPrep normalisation plates (Invitrogen), sequencing was carried out on the Illumina MiSeq platform (600 cycles, V3 chemistry, 300 bp paired end reads) with a 30% PhiX spike (Macrogen).

**Figure 1.**
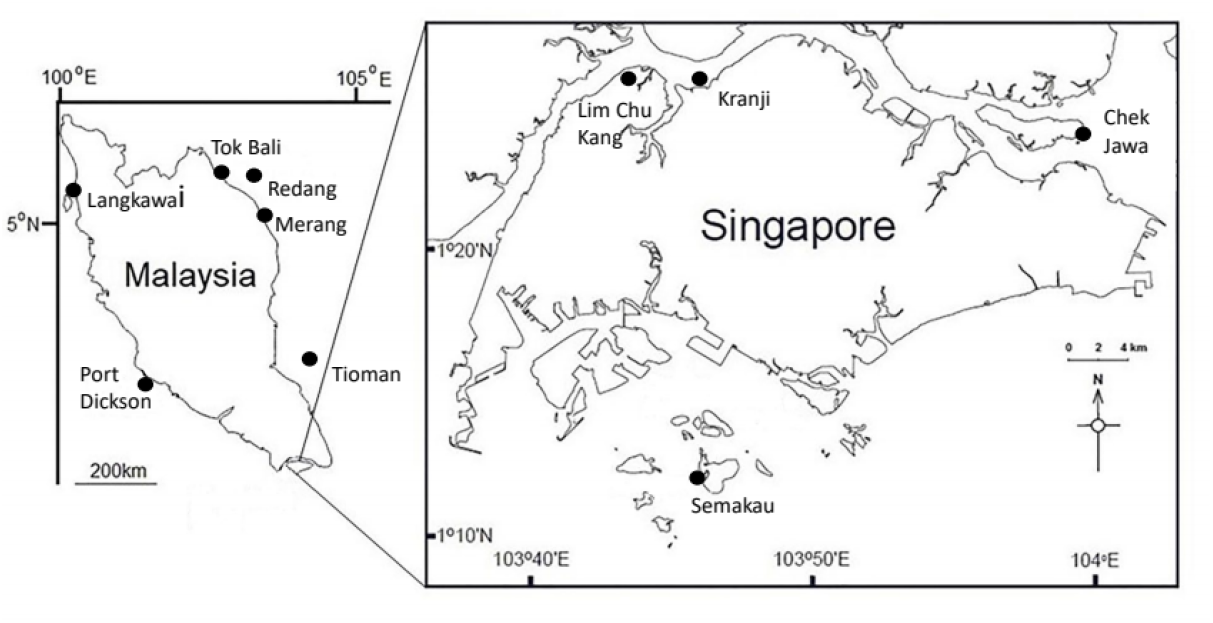
Map indicating sampling locations.

### Bioinformatics and statistics

For full details of sequencing statistics and numbers of reads remaining after quality control and filtering see SI Tables 1&2. Barcodes and adaptors were removed from de-multiplexed sequence files using Cutadapt [65]. Reads were filtered based on quality scores and trimmed using the *DADA2* package version 1.9.0 [66, 67] in R version 3.4.1 (R Core Team, 2017). Forward reads were truncated at 300 base pairs, and reverse reads were truncated at 200 base pairs. Both forward and reverse reads were filtered to remove any reads less than 100 nucleotides long, or with a max EE (expected error) of 2, and reads were additionally truncated at the end of ‘a good quality sequence’ with the parameter truncQ = 2. The DADA2 algorithm was next used to estimate error rates from all quality-filtered reads and then to merge forward and reverse reads and infer amplicon sequence variants (ASVs). Chimeras were removed with de novo detection. Sequenced extraction and PCR negatives were used to identify possible contaminants using the *decontam* R package [67], and remaining ASVs were assigned taxonomy with the RDP classifier [68] against a training set based on the Silva v132 16S database [69]. Phylogenetic placement of ASVs was assigned by aligning sequence variants without an anchor using the AlignSeqs() function of the *DECIPHER* R package version 2.6.0 [70] and constructing a maximum likelihood tree with the optim.pml() function from an initial starting tree built using the NJ() function in the *phangorn* R package version 2.4.0 [71].

Any ASVs assigned to mitochondrial or chloroplast genomes were removed. Raw sequence counts were then converted to relative abundance. Alpha diversity metrics for each site were calculated, and non-metric multi-dimensional scaling (NMDS) was performed on the UniFrac [72] dissimilarity matrix of samples using the *vegan* [73] and *phyloseq* [74] R packages (Fig. 2). Permutational multivariate ANOVA (SI Table 3) was performed on the ASV table with ‘location’ and ‘plant structure’ as predictors using the adonis() function of the *vegan* package. A mantel test with 999 permutations was performed using the vegan R package. Differential abundance analyses were performed using the *vegan, indicspecies* [75] and *corncob* [76] R packages. ASVs and species-level taxa that were detected by all three methods as being differentially abundant in various plant structures are reported below. The *microbiome* R package version 1.10.0 was used to detect a core taxa [77] (Fig. 3). Here we define the core taxa as those present in at least 20% of the samples at relative abundance of at least 10% in each of those samples (i.e., those taxa making up at least 10% of the reads in at least 20% of all samples). All sequences associated with this work have been deposited at the National Center for Biotechnology Information under BioProject ID: PRJNA735404.

**Figure 2.**
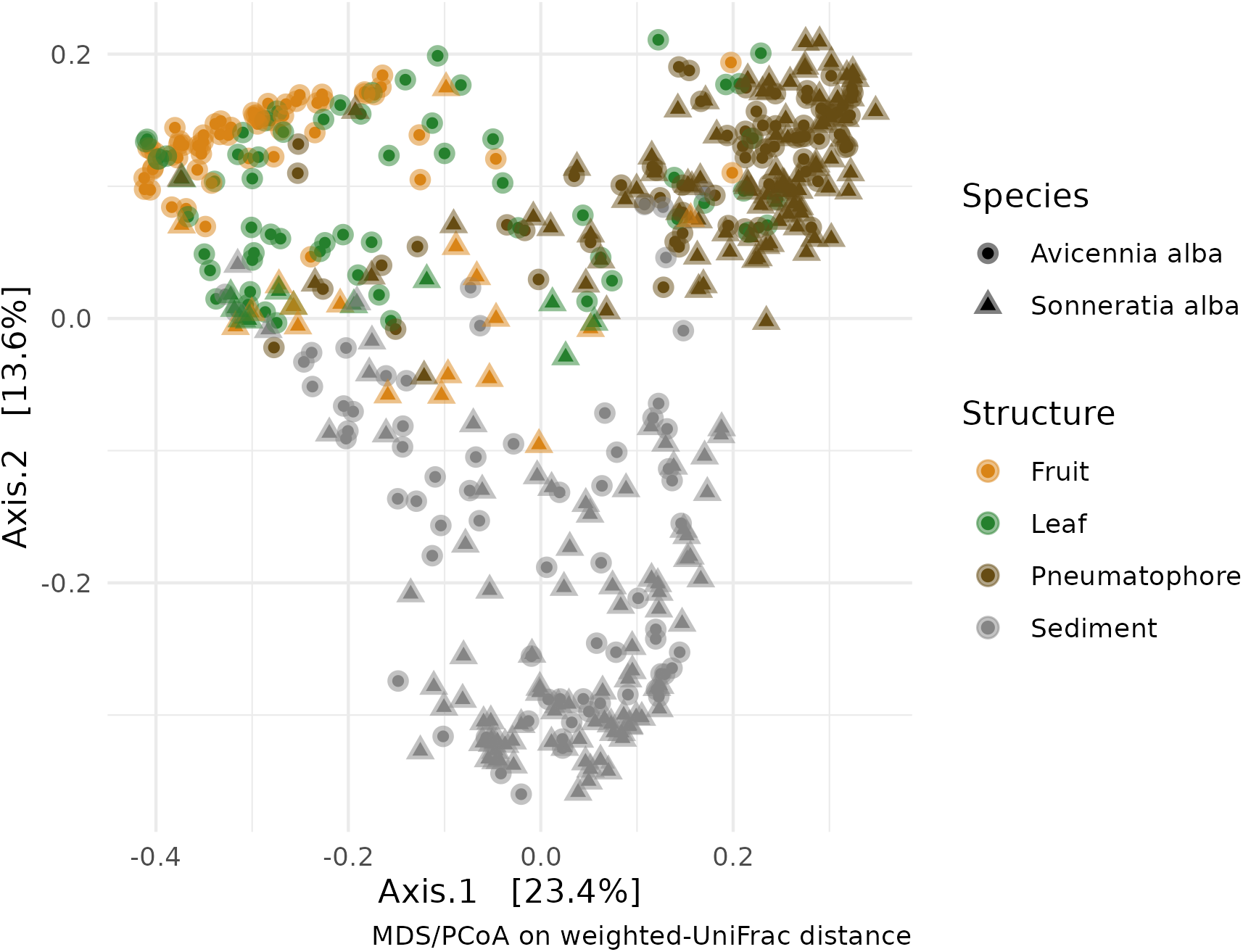
Weighted-UniFrac distance ordination for both species and all sampled parts, including sediment.

**Figure 3.**
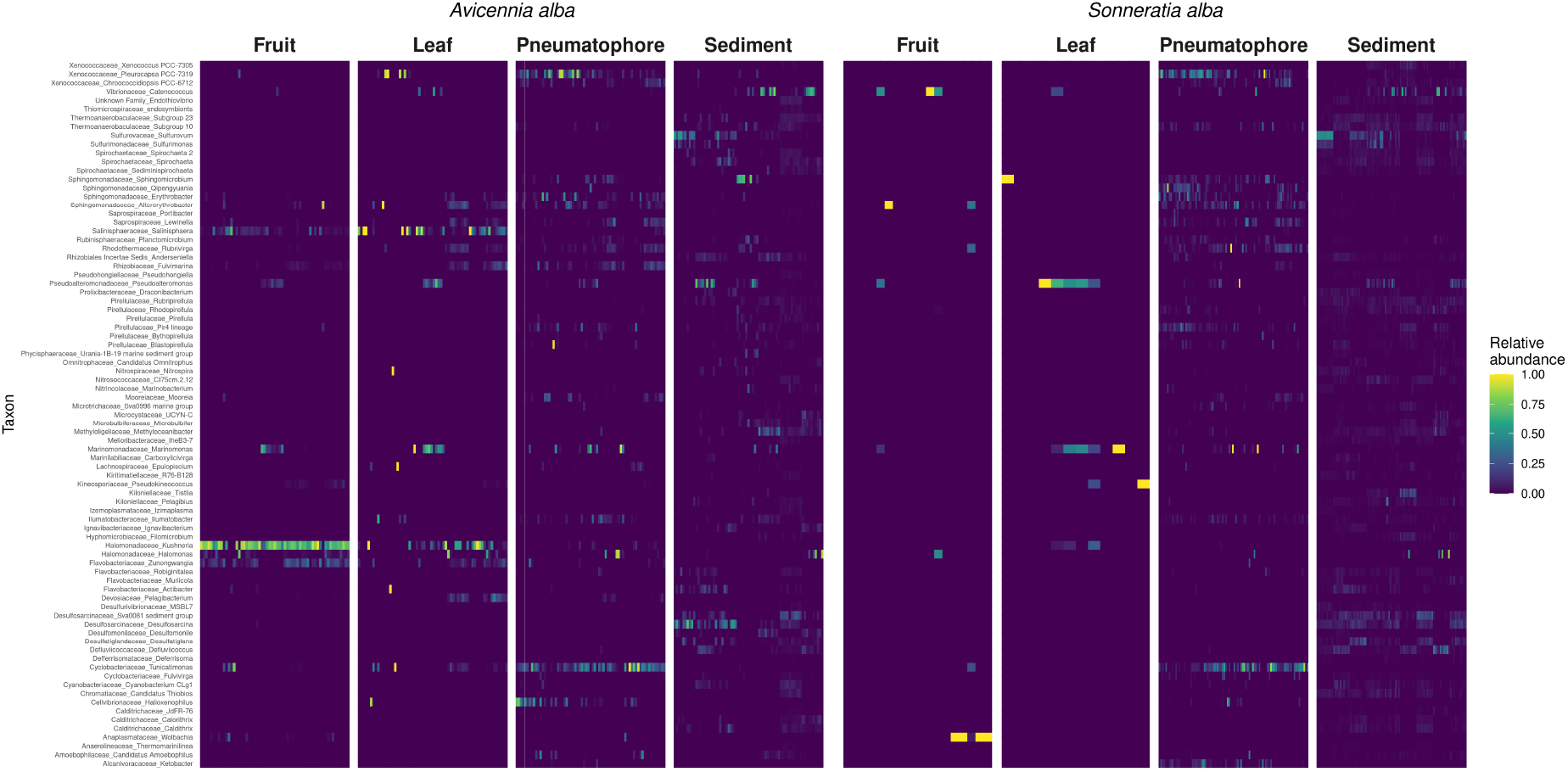
Heatmap showing the abundance of each of the 81 core taxa identified in each species and sampled part.

### Results

In total, sequencing generated 28,297,986 and 29,484,815 reads for *Anicennia alba* and *Sonneratia alba* respectively.After quality control processing and chimera removal, 4,835,163 reads for *A. alba* and 13,861,503 reads for *S. alba* remained and were used in all downstream analysis (SI Tables 1 & 2). Rarefaction curves for both species (SI Fig. 1) indicate that sufficient sequencing depth was achieved, with all samples reaching asymptote.

Permutational multivariate analysis of variance (PERMANOVA) (SI Table 3) indicates a weak but significant difference in bacterial communities between host species (*p* = 0.001, *R*^2^ = 0.015), with sampled part and sampling location having a larger influence on community structure (*p* = 0.001, *R*^2^ = 0.197; & *p* = 0.001, *R*^2^ = 0.097 respectively). This pattern is also evident in the NMDS plot (Fig. 2) with samples of the same type (e.g., leaves) tending to cluster together irrespective of the host species. Above ground parts (leaves and fruits) tend to be similar while sediment and pneumatophores host bacterial communities that are distinct from each other, the leaves, and the fruits. Further confirming this similarity, plots of beta-dispersion indicate a high degree of overlap in microbial communities between species (SI Fig 2) and between fruits and leaves, while pneumatophores and sediment samples tend to be more dissimilar (SI Fig 3).

Differential abundance analysis was performed using Similarity Percentages (SIMPER) in *vegan, indicspecies* and *corncob*. Eight bacterial taxa were identified as differentially abundant between plant components by all three methods. These differentially abundant taxa were *Pleurocapsa, Tunicatimonas, Halomonas, Marinomonas, Rubrivirga, Altererythrobacte, Lewinella*, and *Erythrobacter* (Fig 4.). The highest bacterial diversity was found in sediment samples. In living tissues in both mangrove species, pneumatophores contained the highest diversity, while leaves and fruits showed similar diversity (SI Fig. 4). A core microbiome consisting of 81 taxa was identified in both species throughout all samples (Fig. 3). A mantel test indicated a significant pattern of distance decay of similarity in fungal communities assessed (999 permutations; Mantel statistic r: 0.1097; *p* = 0.001)

**Figure 4.**
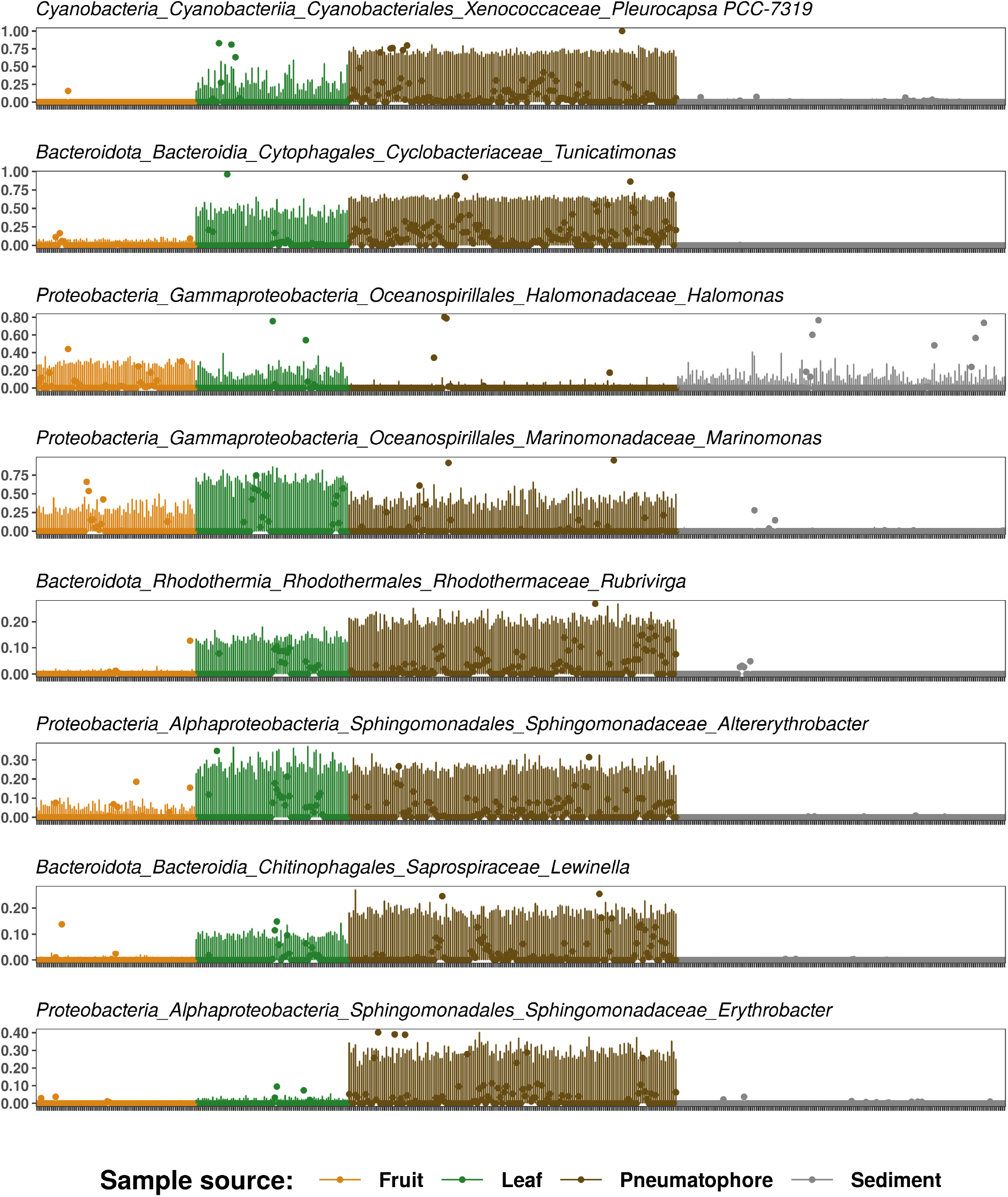
Plot indicating differential abundance of each of the eight taxa identified using three different techniques.

## Discussion

Consistent with other work examining microbial community structure in a variety of host taxa, we show that microbial community structure does exist around the Malay Peninsula [39, 40, 54, 55, 78, 79], and this structure is driven primarily by plant part sampled, location sampled, and host identity. It is likely that these differences are driven by local environmental conditions, with locations closer together tending to have more similar environmental conditions in comparison to those that are more geographical distant. This idea is further supported by the significant pattern of distance decay of similarity we observe in microbial communities, meaning that communities are more similar when spatial distance between them is low, a likely consequence of the host’s ability to exert a degree of control over the composition of their microbial communities, tailoring them to local conditions [80, 81]. Given these findings, and the recognised high rates of failure in mangrove restoration programmes, a consideration of microbial communities could help to improve success [82], more so since inoculation with bacterial and fungal species found in the local environment has already been shown to improve restoration success [5, 60].

We identified a core *Anicennia alba* and *Sonneratia alba* mangrove microbiome consisting of 81 taxa. Given the ubiquity of these taxa in both species and in all living parts (e.g., leaves and pneumatophores) it is likely they play a key role in maintaining or promoting host fitness. For example, mangroves are nitrogen limited environments, and in general are considered as nutrient-deficient habitats [83–85], correspondingly a number of the taxa we identified in the core microbiome are involved in nitrogen cycling and thus help promote growth in what would otherwise be a challenging environment for plant life. Similarly, the core microbiome in this work contains a number of taxa that are involved in sulphur cycling. The bacterial oxidation of sulphur can improve substrate fertility and aid in the removal of toxic sulphide that is produced by sulphur reducing bacteria [86, 87]. These core taxa would be a good starting point for further investigation as potential inoculates that could improve restoration success, especially when transplants are grown *ex-situ*. However, increasingly evidence is also suggesting that rare taxa are likely to be just as important [62] and may play important roles in allowing hosts to survive in challenging environments, or in helping facilitate adaption to geographically unique environmental conditions. This is especially true in biogeochemical cycles and in protecting hosts from pathogens [88, 89].

We identified eight bacterial taxa shared between both species that are differentially abundant (Fig. 4). These eight were primarily enriched in living parts suggesting that they play a vital role in the Southeast Asian mangrove microbiome. *Pleurocapsa*, a nitrogen fixing cyanobacteria [90] is enriched within pneumatophores, and it is likely that members of this group facilitate nitrogen uptake in the water logged and nutrient limited mangrove sediments. Members of the Bacteroidetes genus *Tunicatimonas* (here, enriched in pneumatophores) were first isolated from sea anemones [91] and have since been identified as common plastic debris in marine environments, where they are thought to be well adapted to take advantage of the new niches this plastic creates [92]. While *Tunicatimonas* might not serve any purpose in promoting host health, their increased abundance could be a consequence of plastic pollution. Plastic pollution is a recognised problem in Southeast Asia, and much of this plastic can become entangled within mangroves where it can disrupt growth and lead to ecological instability [93, 94]. We hypothesize that plastic debris could be facilitating the transport of t*Tunicatimonas* into mangrove ecosystems, particularly since *Tunicatimonas* taxa are found enriched on pneumatophores, a structure ideally suited to trapping plastic debris. *Halomonas* spp. (here, enriched in sediment, fruits, and leaves) have previously been identified in Southeast Asian mangroves where they produce the compound ectoine [95]. This compound is able to stabilize proteins and other cellular structures in the presence of high intensity UV irradiation, heat stresses (cold and high), fluctuations in pH and can help hosts survive extreme osmotic stress [95, 96]. Members of this genus are most enriched in the above ground structures - those that are exposed to the extreme levels of UV light that are frequently encountered in tropical habitats suggesting that they could play a protective role in these parts. The genus *Marinomonas (here, enriched in all living plant tissues)* is abundant in marine ecosystems where it is implicated in melanin synthesis and the catabolism of dimethylsulfoniopropionate (DMSP) to dimethyl sulfide (DMS). DMSP metabolisers are acknowledged as important constituents of the coral microbiome where they have principal roles in sulphur cycling, pathogen suppression and mediating thermal stress responses [78, 97–99]. The same properties that make DMSP metabolisers beneficial microorganisms in coral mean that this group warrants further investigation in mangroves, particularly as they are enriched in both species. Little is known of the *Rubrivirga* genus (here, enriched in leaves and pneumatophores), however, they are described as chemoorganotrophs [100] meaning they are able to oxidise organic chemicals to produce energy. Therefore, they are likely to be involved in nutrient cycling and this genus is a good candidate for further investigation into its role in the mangrove microbiome. Similarly, details on *Altererythrobacter* (here, enriched in leaves and pneumatophores) are scarce, with no information on what functions they perform, but they have been isolated from mangrove ecosystems previously [101, 102]. This ubiquity suggests they have an important role in the mangrove microbiome and would be ideal candidates for further study. *Lewinella (here, enriched in leaves and pneumatophores)* are halotolerant heterotrophs that are commonly encountered in activated sludge and are able to break down complex molecules [103]. They show increased abundance when coastal areas undergo sudden vegetation declines that can arise through land use change [104], with further investigation of this genus it may be possible to identify indicator species that can be used in the design of pro-active management strategies. *Erythrobacter (here, enriched in pneumatophores)* have been isolated from mangrove habitats in multiple studies throughout the world [84, 105–107], but little is known about what they are doing in these habitats. Like the *Altererythrobacter*, the ubiquity of *Erythrobacter* in multiple mangrove forests suggests that they play an important role in the mangrove microbiome, and again this genus would be a worthy candidate for future studies trying to determine whether beneficial mangrove microbes exist.

Despite having a shared core of 81 taxa across all living parts, each part does host a significantly different bacterial community. In both species, leaves and fruiting bodies, or above ground structures have a similar diversity, of the living parts the pneumatophore has the most diverse microbial community. Similar to other studies the highest diversity of microbes is seen in sediment samples [39, 40, 58, 59]. Our ordinations further support these observations with fruit and leaves having similar microbial communities and therefore clustering together, while pneumatophores and sediment samples form well-defined and clearly differentiated clusters. In all cases, the microbial communities from the same species, in the same sampled part (e.g., *A. alba* and *S. alba* leaves) appear similar and do not form distinct clusters, this similarity between species corroborates the high number of shared core taxa between both species.

Microbes have fundamental roles in the cycling of many elements, and are major regulators of greenhouse gasses [108]. Climate change will alter how these microbial communities are structured [17, 19, 22, 24], and experimental manipulations show that 4°C of warming can increase soil respiration by as much as 37%, with much of this increase mediated by microbial decomposition of sequestered carbon [22]. Under the anticipated future climate change scenarios, much of the carbon stored in coastal ecosystems will become vulnerable to increased microbial decomposition, and mangrove ecosystems will likely move from net sinks to net sources of greenhouse gasses. Given this, it is crucial that we understand how mangrove microbial communities are currently assembled and structured in order to determine how best to keep existing carbon stores intact.

The arguments in favour of mangrove restoration to increase biodiversity are unequivocal. However, things are much less clear when it comes to blue carbon and how climate change and restoration will impact greenhouse gas fluxes in coastal ecosystems. Anecdotally, the consensus suggests that restoration will aid in the sequestering of atmospheric carbon. Unfortunately, scientific surveys of greenhouse gas fluxes in restored ecosystems shows that CO_2_ fluxes in these ecosystems is actually increased. These post restoration increases are observed in numerous coastal ecosystems (e.g., mangroves, kelp forests, saltmarshes and seagrasses) [28, 104, 109–112] and demonstrate the complexities involved when trying to understand greenhouse gas fluxes in these ecosystems. If all factors (e.g., microbial communities) are not considered, it is possible that we will do more harm than good, particularly when it comes to using coastal ecosystems in carbon trading schemes and in the offsetting of emissions.

This work uncovers a core mangrove microbiome and identifies a more limited set of eight microbial taxa that are differentially abundant in certain plant components in both species of mangroves surveyed. These would be good candidate taxa for further screening since their presence in both host species and their elevated abundance suggests possible beneficial properties that could improve restoration success. Importantly, this work also characterizes the microbes in the sediment where the majority of the sequestered carbon is stored, providing a baseline for future studies of how climate change is impacting this crucial ecosystem.

## Author Contribution

Study conception and design, BJW. Sample collection, BJW. Laboratory work BJW. Analysis, BJW, TM, LB, KH, ZYY, GZ. First draft, BJW, GZ. All authors commented and contributed to improving the manuscript and preparing the manuscript for submission. All authors read and approved the final manuscript.

## Data Availability

Raw sequences obtained in this study have been deposited in the National Center for Biotechnology Information under BioProject ID: PRJNA735404

